# PanoView: An iterative clustering for single-cell RNA sequencing data

**DOI:** 10.1101/616862

**Authors:** Ming-Wen Hu, Dong Won Kim, Sheng Liu, Donald J Zack, Seth Blackshaw, Jiang Qian

## Abstract

Single-cell RNA-sequencing (scRNA-seq) provides new opportunities to gain a mechanistic understanding of many biological processes. Current approaches for single cell clustering are often sensitive to the input parameters and have difficulty dealing with cell types with different densities. Here, we present Panoramic View (PanoView), an iterative method integrated with a novel density-based clustering, Ordering Local Maximum by Convex hull (OLMC), that uses a heuristic approach to estimate the required parameters based on the input data structures. In each iteration, PanoView will identify the most confident cell clusters and repeat the clustering with the remaining cells in a new PCA space. Without adjusting any parameter in PanoView, we demonstrated that PanoView was able to detect major and rare cell types simultaneously and outperformed other existing methods in both simulated datasets and published single-cell RNA-sequencing datasets. Finally, we conducted scRNA-Seq analysis of embryonic mouse hypothalamus, and PanoView was able to reveal known cell types and several rare cell subpopulations.

**Author summary:** One of the important tasks in analyzing single-cell transcriptomics data is to classify cell subpopulations. Most computational methods require users to input parameters and sometimes the proper parameters are not intuitive to users. Hence, a robust but easy-to-use method is of great interest. We proposed PanoView algorithm that utilizes an iterative approach to search cell clusters in an evolving three-dimension PCA space. The goal is to identify the cell cluster with the most confidence in each iteration and repeat the clustering algorithm with the remaining cells in a new PCA space. To cluster cells in a given PCA space, we also developed OLMC clustering to deal with clusters with varying densities. We examined the performance of PanoView in comparison to other existing methods using ten published single-cell datasets and simulated datasets as the ground truth. The results showed that PanoView is an easy-to-use and reliable tool and can be applied to diverse types of single-cell RNA-sequencing datasets.

## Introduction

Single-cell RNA-sequencing (scRNA-seq) has attracted great attention in recent years. Unlike traditional bulk RNA-seq analysis, scRNA-seq provides access to cell-to-cell variability at the single-cell level. This allows defining individual cell types, and subtypes, among a population containing multiple types of cells, and also makes possible following how individual cell types change over time or after being exposed to various perturbations (1–4).

Classifying single cells based on their expression profile similarity is the basis for scRNA-seq analysis. A variety of clustering approaches have been developed and applied to scRNA-seq analysis such as hierarchical clustering (5–7), K-means clustering(8–11), SNN-Cliq(12), pcaReduce(13), SC3(14), Seurat(3,15), SCANPY(16), RCA(17), and dropClust(18). There are also algorithms, like RaceID/RaceID2(4,19) and GiniClust (20), were developed specifically to identify rare cell types. Nevertheless, one challenge is that clustering results are often highly sensitive to input parameters, and sometimes the required parameters are not intuitive to users (S1 Table). For example, DBSCAN(21) is a clustering that required two parameters to classify clusters based on the densities of subpopulations, and has been applied in some scRNA-seq studies(3,22,23). However, it is difficult for users to pick proper required parameters without the aid of other computer programs and different parameters can lead to different clustering results (S1 Fig and S2 Fig). Furthermore, it is also challenging for density-based clustering algorithms to properly handle clusters with different densities(23). This can often be the case for single cell clustering because different cell types can exhibit different levels of variation in similarity among the cluster members.

To address these issues, we have developed Panoramic View (PanoView), which utilizes an iterative approach that searches cell types in an evolving principal component analysis (PCA) space. The strategy is that we identify the cell cluster with the most confidence in each iteration and repeat the clustering algorithm with the remaining cells in a new PCA space (Fig 1A). We define the most confident cluster as the “mature” subpopulation that has the lowest variance in the current PCA space. To cluster cells in a given PCA space, we have developed a novel density-based algorithm, namely Ordering Local Maximum by Convex hull (OLMC) (Fig 1B-D), that uses a heuristic approach to estimate the required parameters based on the input data structures (see Methods).

**Fig 1.**
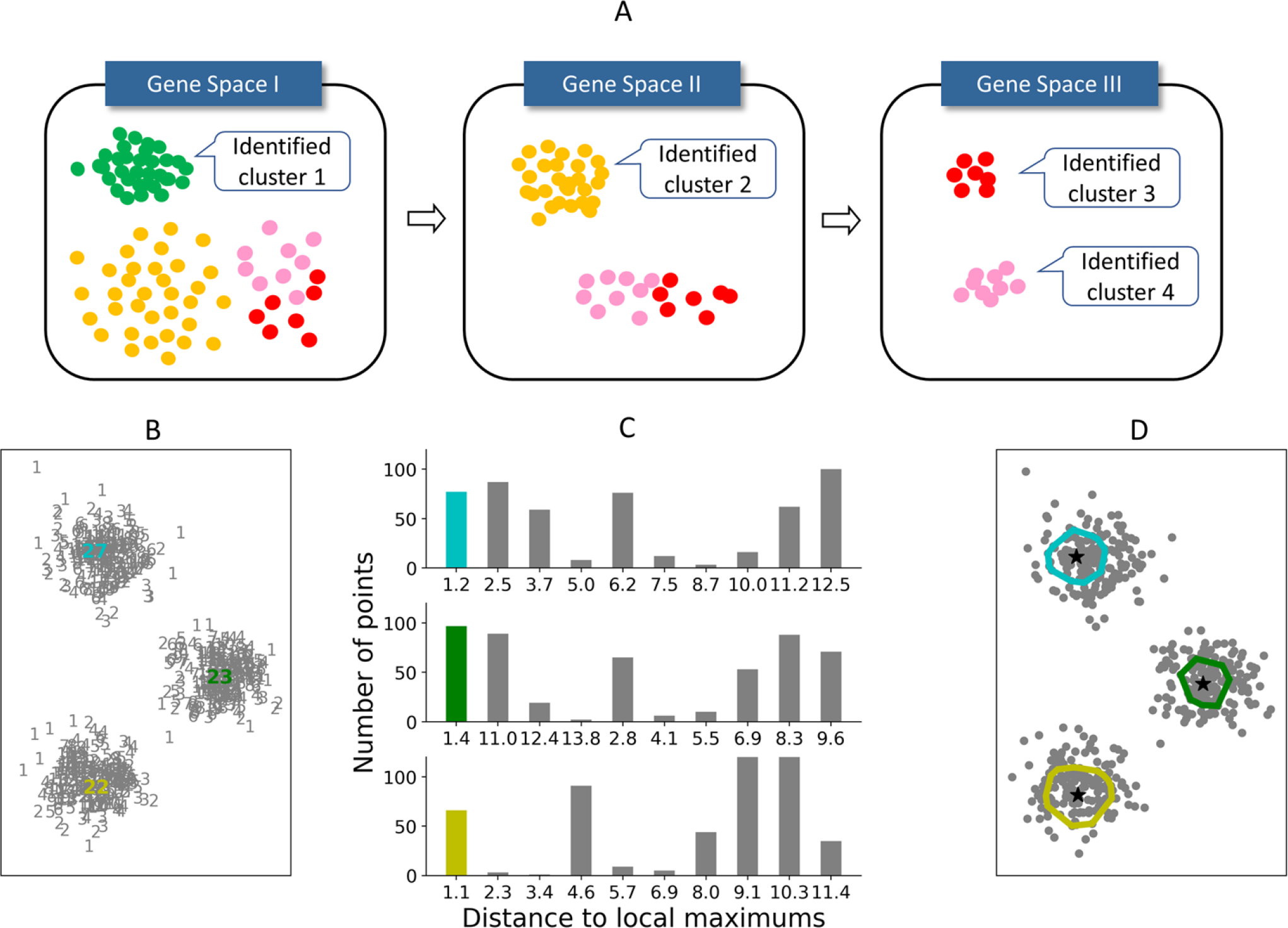
Panoramic View algorithm. (A) The schematic illustration of PanoView algorithm. (B-D) A toy model for the illustration of OLMC algorithm. (B) 500 random points in 2D space. Gray numbers represent the number of neighbors for each point. Colored numbers are three local maximum densities. (C) The histograms represent the distance to local maximums. The heights of colored bars are used for constructing the first convex hull for each local maximum. (D) Color-enclosed circles represent the convex hulls constructed by colored bars in (C) during OLMC algorithm.

## Results/Discussion

### Results of simulated datasets

To evaluate the performance of PanoView, we first tested 1,200 simulated data with varying configuration parameters (e.g. numbers of clusters and standard deviation of the members within clusters). The performance of the clustering was evaluated using the Adjusted Rand Index (ARI), which measures the similarity between the cell membership produced by a chosen method and the ground truth(24).

We compared the performance of PanoView with 9 existing methods, including pcaReduce(13), SC3(14), Seurat(15), SCANPY(16), RCA(17), K-means without prior dimensional reduction, PCA followed by DBSCAN, PCA followed by K-means, and TSNE followed by K-means. The results showed that PanoView and SCANPY outperformed other benchmarking methods in all datasets tested using default parameters. Although we input the correct number of clusters for K-means and pcaReduce, their performance decreased in the datasets with a large number of clusters (K-means, TSNE+Km, PCA+Km, pcaReduce in Fig 2A). For DBSCAN, we tuned the required parameters until they reached optimal performance in datasets with *n*=3 and 4 (PCA+DB in Fig 2A). However, its performance dropped significantly when *n* > 10. We also observed a similar outcome in Seurat, whose performance dramatically dropped for *n* > 17. It is worthy to note that these methods could achieve much better performance if we tune the parameters for each dataset. In this study, we only used the default parameters for all the methods and evaluated the robustness of the methods with different datasets. SC3 and RCA with default parameters did not produce usable clustering result for the simulated datasets.

**Fig 2.**
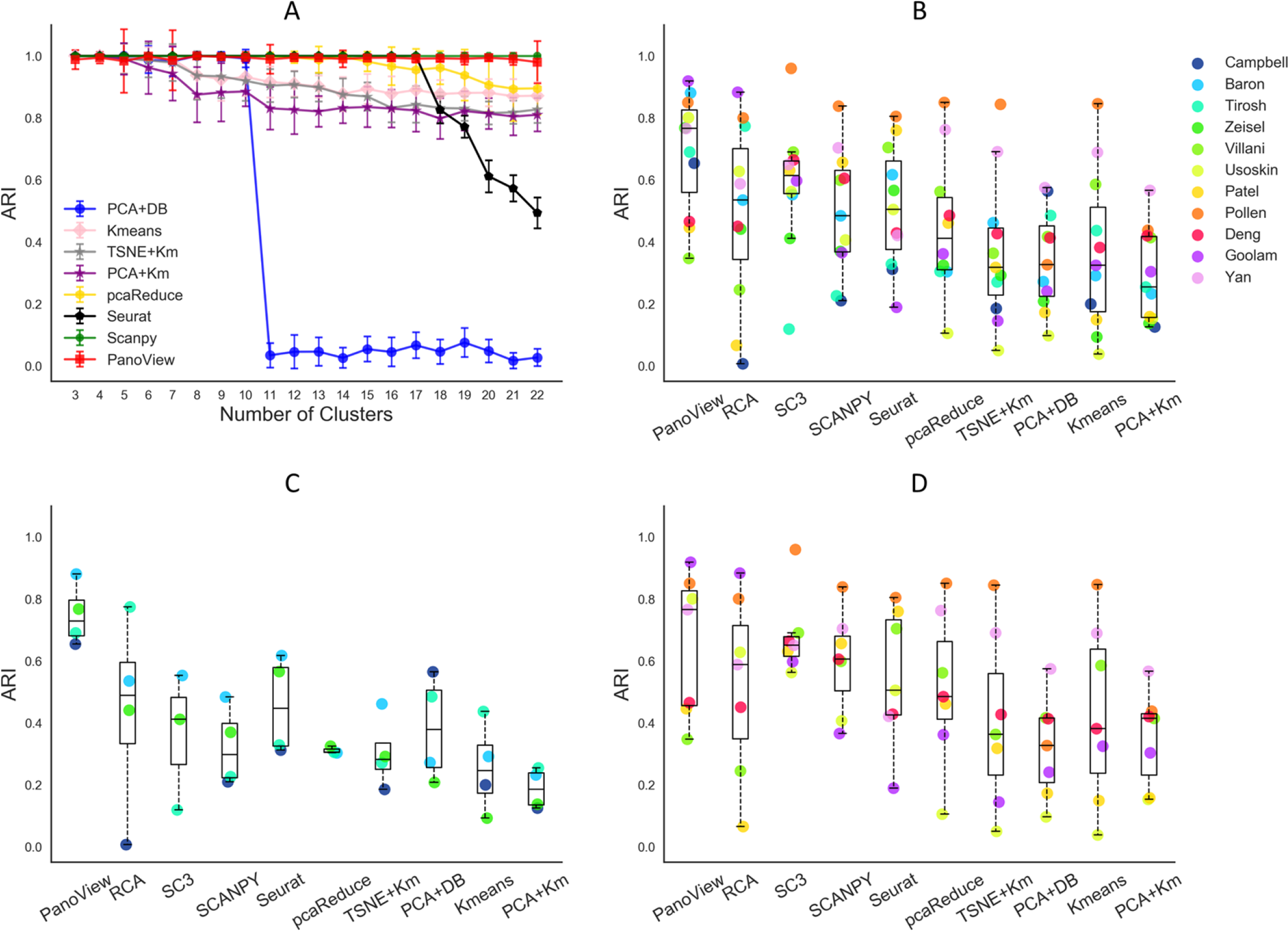
The performance of PanoView in comparison to other existing methods using ten simulated datasets and published scRNA-seq datasets. (A) The ARI results of 8 different computational methods in 1,200 simulated datasets. Error bars indicate standard deviation of ARI. (B) The ARI result of 10 clustering methods in 11 published single-cell RNA-seq datasets. The order of legend is based on the number of single-cells in descending order. Dots are the calculated ARI values for each dataset. SC3 and pcaReduce did not produce usable clustering results for Campbell dataset. The dataset is missing for these two methods in B and C. (C) The ARI result of 4 datasets that contain more than 3,000 cells. (D): The ARI result of 7 datasets that contain fewer than 3,000 cells.

### Results of published scRNA-seq datasets

We applied PanoView to 11 published scRNA-seq datasets, ranging in size from 90 cells to 20,921 cells (S2 Table). We used the reported clustering results as the ground truth for the calculation of ARI, assuming that the authors optimized their analysis correctly with the expertise in the research topics. Based on the overall performance of eight tested methods, we divided them into two tiers by the median value of 0.5 in ARI (Fig 2B). The median values of ARI in the first tier are 0.766 (PanoView), 0.614 (SC3), 0.535 (RCA), and 0.505 (Seurat). For the second tier, the median values are 0.483 (SCANPY), 0.411 (pcaRecue), 0.327 (PCA+DB), 0.325 (Kmeans), 0.255 (PCA+Km), 0.318 (TSNE+Km). This difference in tiers was not surprising, as the methods in the first tier were specifically designed for single-cell analysis. Though SCANPY and pcaReduce were also developed for the analysis of single cells, they did not show good performance in this study. In the first tier, four methods seem to have relatively similar performance. However, there is a noticeable difference in the datasets that exceed 3,000 cells. Fig 2C shows that for these larger datasets, PanoView outperformed the other methods by a significant margin, so that the median value of ARI was 0.729 and the rest of methods were 0.488 (RCA), 0.411 (SC3), 0.298 (SCANPY), 0.447 (Seurat), 0.305 (pcaReduce), 0.282 (TSNE+Km), 0.378 (PCA+DB), 0.245 (Kmeans), 0.185 (PCA+Km). We also observed that PanoView displayed relatively less variation. For smaller datasets, PanoView (median: 0.766) still ranked first among all methods tested (Fig 2D). The result of ARI values for all methods is provided in S3 Table.

### Computational cost

We also examined the computational cost of PanoView in the real scRNA-seq datasets. It is not surprising that data analysis takes longer when datasets contain more cells (Fig. 3). We also compared the computational cost with other methods, which generated reasonable clustering results. It is obvious that PanoView is not the fastest algorithm. SCANPY, Seurat and RCA are faster than PanoView. It is interesting that SC3 and pcaReduce are slower than PanoView and they failed to generate clustering results for the largest dataset.

**Fig 3:**
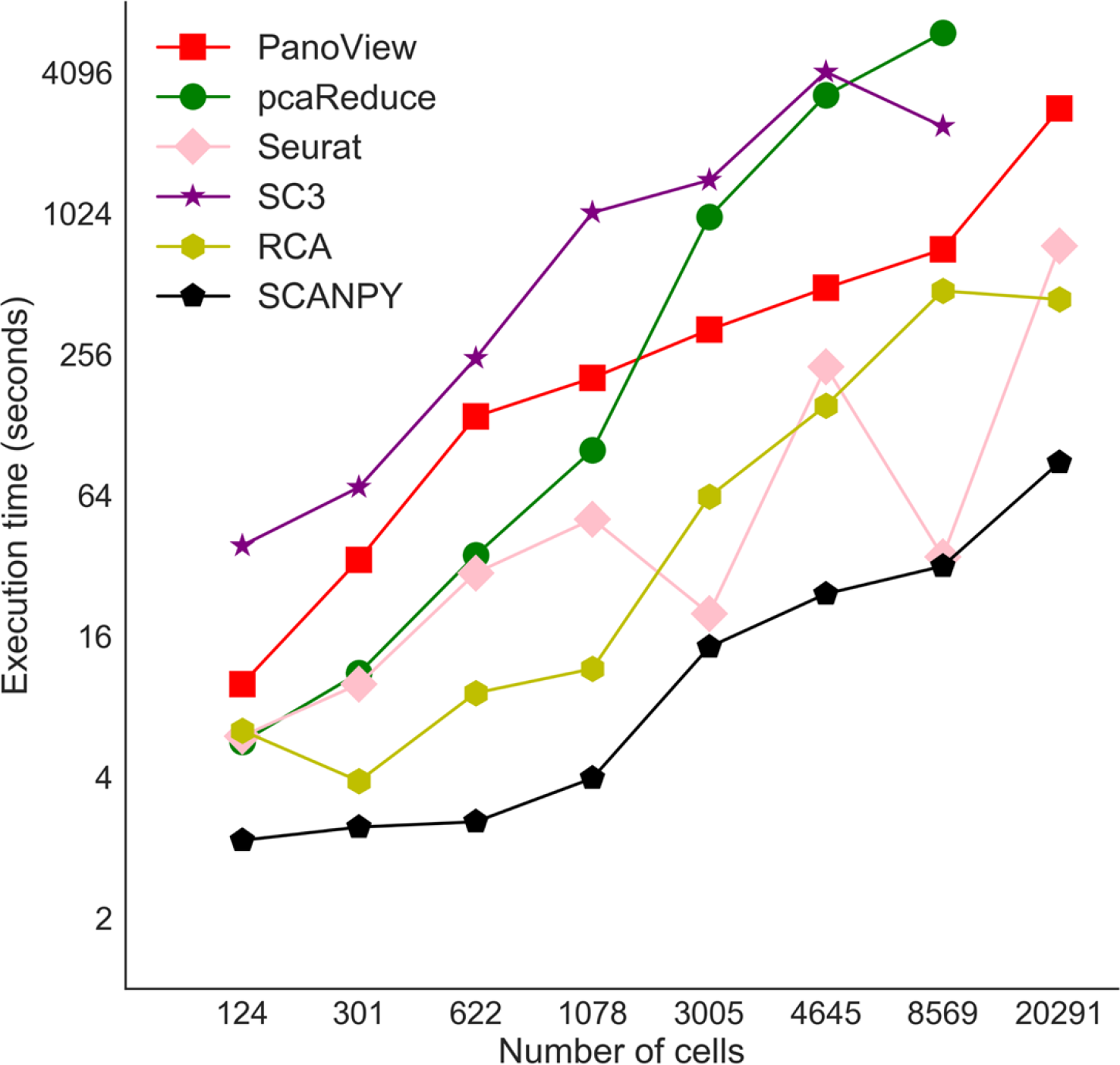
Computational times for selected clustering methods. The X-axis represents the number of cells in 8 datasets. Note that SC3 and pcaReduce did not produce usable clustering result for Campbell dataset.

### Stability of default parameters in PanoView

PannoView produced the results using default parameters. We investigated whether the default parameters produced the optimal results. We have 8 variables in the PanoView, including Zscore, Gini, Bc, Bg, Maxbb, CellNumber, GeneLow, Fclust_height (see Methods for details). If we provided 3 values for each variable, including one default value, we have 6,561 different combinations of these parameters. Since it takes too long to use all potential combinations, we executed PanoView with 500 randomly picked combinations on the 10 scRNA-seq datasets (To save the computational time, we didn’t include the Campbell et al dataset for this analysis). Based on the sampling results from the 500 combinations, we found that the default parameter set could produce overall good clustering results across the 10 datasets, ranking in the top 98.2 percentile among the 500 parameter sets (Fig 4A-B). The similar observation was made for each individual dataset (Fig. 4C), although the default parameters performed better in some datasets than others. The 500 clustering results are provided in Table S4.

**Fig 4.**
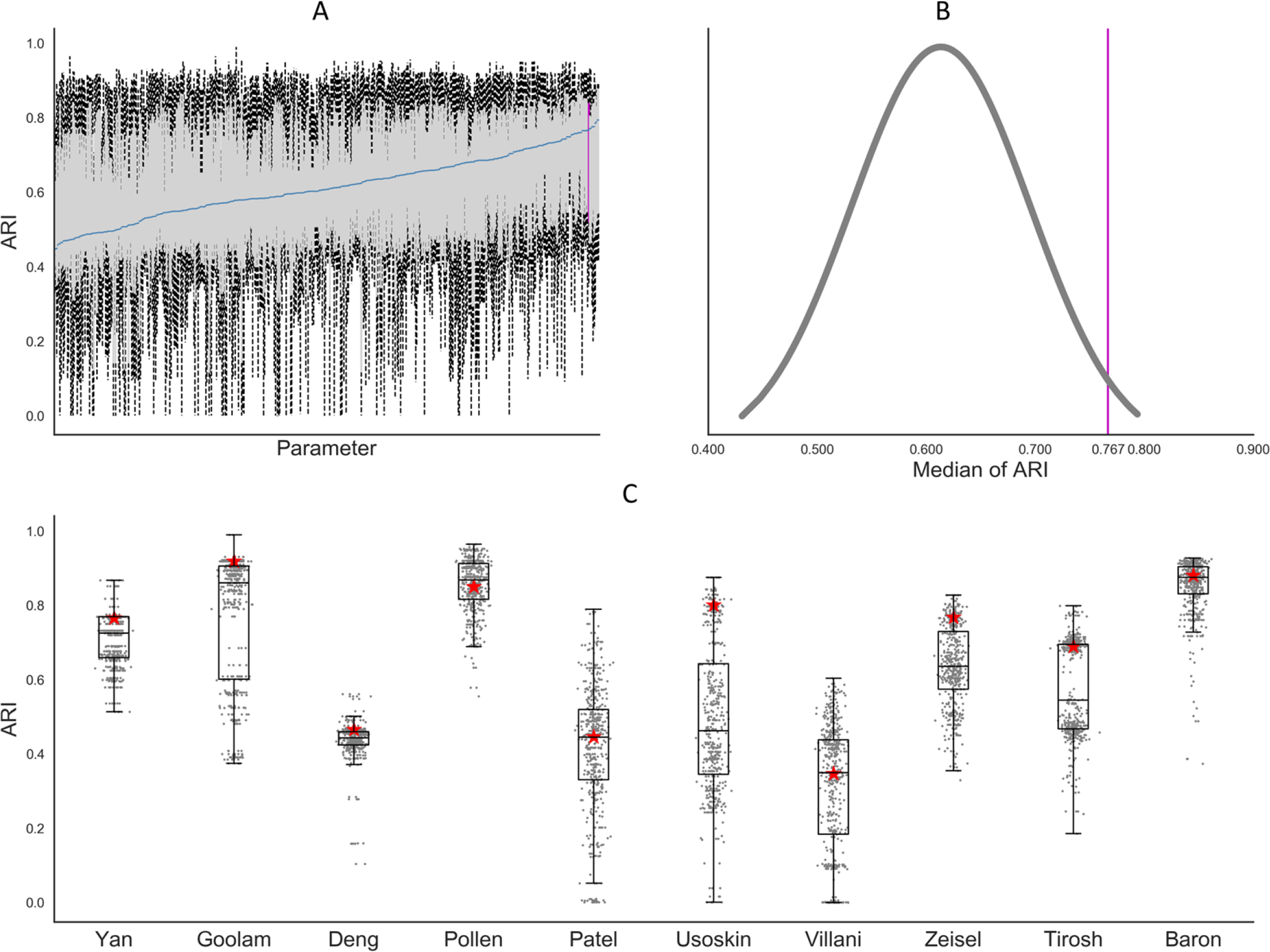
The results of PanoView for 10 scRNA-seq datasets with 500 random parameter sets. (A) Boxplots of 500 simulation results. We ordered the 500 parameter sets based on median values of ARI in ascending order. The blue line indicates the median values of 10 ARI values for each parameter set. The vertical pink line represents the result of PanoView with the default parameters. (B) The distribution of median ARI in 500 simulation results. The default value ranked 98.2 percentile. (C) Boxplots of 500 clustering results in 10 scRNA-seq datasets. Red stars are the ARI results with current default parameters.

### Results of detection of rare cell types

To evaluate the ability to identify rare cell types, we first applied PanoView to 260 simulated datasets and benchmarked it with Seurat, GiniClust, RaceID2, and SCANPY. GiniClust and RaceID2 are two single-cell methods that were specifically designed for detecting rare cell types. We used recovery rate and false positive rate to evaluate the performance of detecting rare cell types (table in Fig 5). PanoView had the best performance that it correctly recovered the rare cell subpopulation in 87.31% of datasets. Although GiniClust recovered 66.54% of datasets, there were 85 datasets contained false-positive rare clusters, resulting in a false-positive rate of 32.69%. In the case of PanoView, only 6 datasets had false-positive rare clusters, resulting in a false-positive rate of 2.3%. Seurat had 3 false-positive rare clusters, resulting in a false-positive rate of 1.15%. We used one simulated dataset to illustrate the accuracy between methods (Fig 4B–4F). PanoView is the only method that perfectly identified rare cell populations and major cell populations. GiniClust did recover the rare cell populations; however, it also produced false positive cells that were scattered in the three other major clusters. Seurat and SCANPY also showed poor performance in identifying rare cell types. Specifically, Seurat divided the one rare cell type into three clusters, while SCANPY grouped rare cells into one major cluster. RaceID2 did not produce a usable clustering result for this chosen dataset.

**Fig 5.**
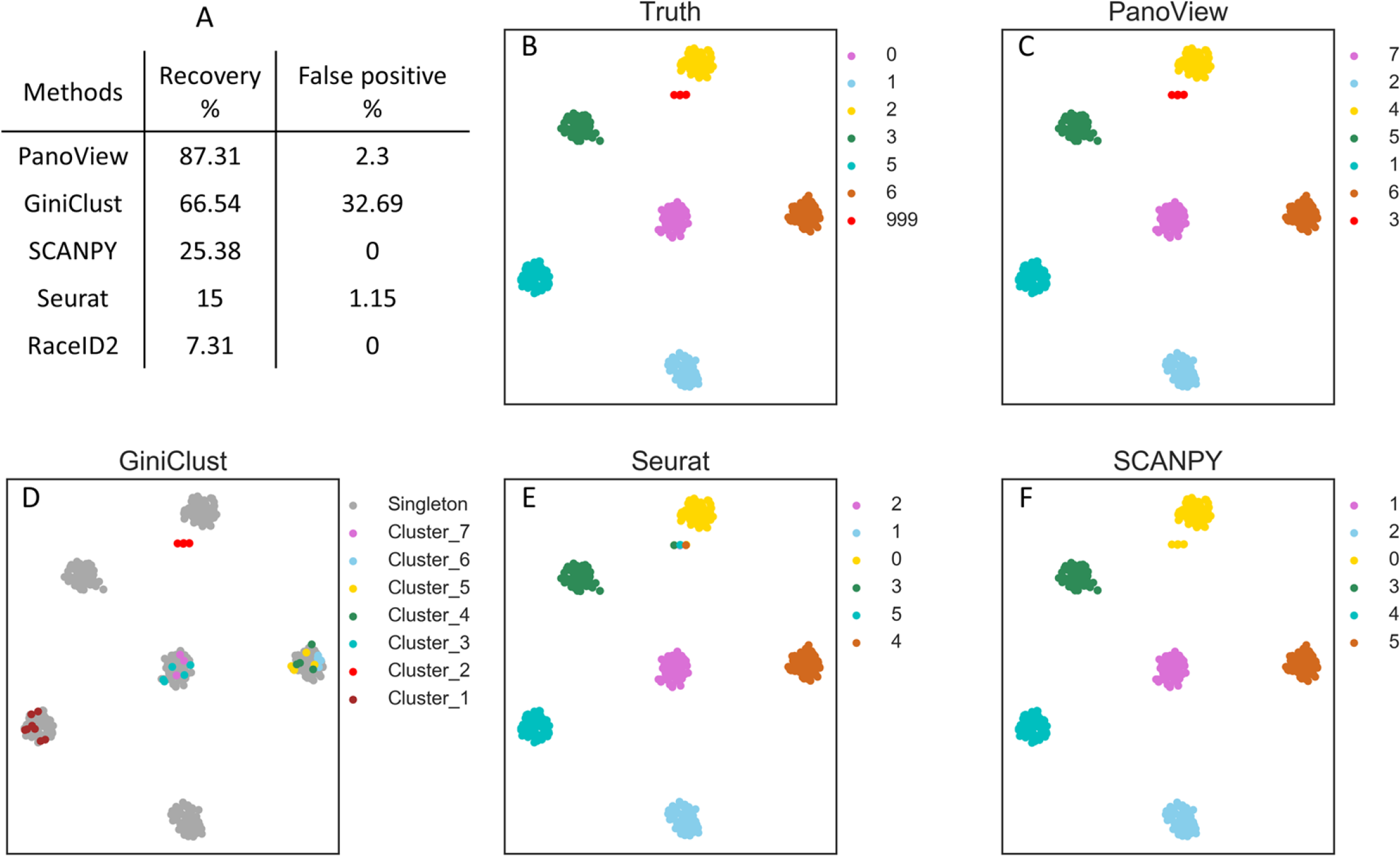
The evaluation of detecting rare cell types. (A) The recovery rate and false positive rate in detecting rare cell types in 260 simulated datasets. SC3 was not included in the comparison because it did not produce usable results in our simulation (B) The ground truth of the selected simulated data. Cluster 999 represents the predefined rare cell type and the TSNE coordinates of three rare cells were adjusted for better visualization. (C, D, E, F) We selected one of the simulated datasets to visualize the performance of different computational methods (PanoView, GiniClust, Seurat, SCANPY). RaceID2 did not produce clustering result in this simulated dataset.

In addition to simulated datasets, we also used Patel dataset to examine the performance of detecting rare cells (Fig. S4). GiniClust reported that it successfully detected one rare cell type in this dataset (20), which consists of 9 cells in glioblastoma tumors. These cells were also discovered by the original study showing highly expressed oligodendrocyte genes(6). In our result (Fig S4), PanoView identified a cluster (cluster #2) that includes 7 cells, which are corresponding to the rare cells in the original study. SCANPY reported a cluster with 9 cells, among which 8 were the rare cells. SC3 identified a cluster with 10 cells, among which 8 were the rare cells. Seurat assigned 9 rare cells to a major cluster, which has 88 cells in total. A similar outcome was also observed in RCA and pcaReduce that both algorithms merged the rare cells to a major cluster. RaceID2 recovered 8 rare cells from a cluster with 9 cells; however, it also produced many much smaller clusters than the other methods. These results indicated that PanoView not only recovers most rare cells but also produces reasonable clusters representing the heterogeneity in glioblastoma tumors cells.

### Clusters of single-cell subpopulations in mouse embryonic hypothalamus

Finally, we applied PanoView to a newly generated scRNA-seq dataset of 959 cells obtained from embryonic day 16 (E16) mouse hypothalamus. The mammalian hypothalamus, which is the central regulator of a broad range of physiological processes and behavioral states, is highly complex at the cellular level (25–27). Cell subtypes in the developing hypothalamus are very poorly characterized. PanoView identified a total of 11 clusters (Fig 6A), the majority of which consisted of radial glia, neurogenic and gliogenic progenitor cells, immature neurons, as expected. A considerable number of non-neuronal cells were also profiled, including pericytes, endothelial cells, erythrocytes, and macrophages. We selected 12 marker genes to show the specific expression level across 11 clusters (Fig. 6B). Four rare cell clusters were also identified, which consisted of a myeloid-like cell type that likely consists of pericyte precursors (28), tissue-resident microglia, infiltrating monocytes, and an unidentified vascular cell type. With the exception of the last cell type, which likely represents a previously uncharacterized subtype of endothelial or pericyte precursor cell, the other three rare cell types represent cells that are known to be found in the embryonic mouse brain.

**Fig 6.**
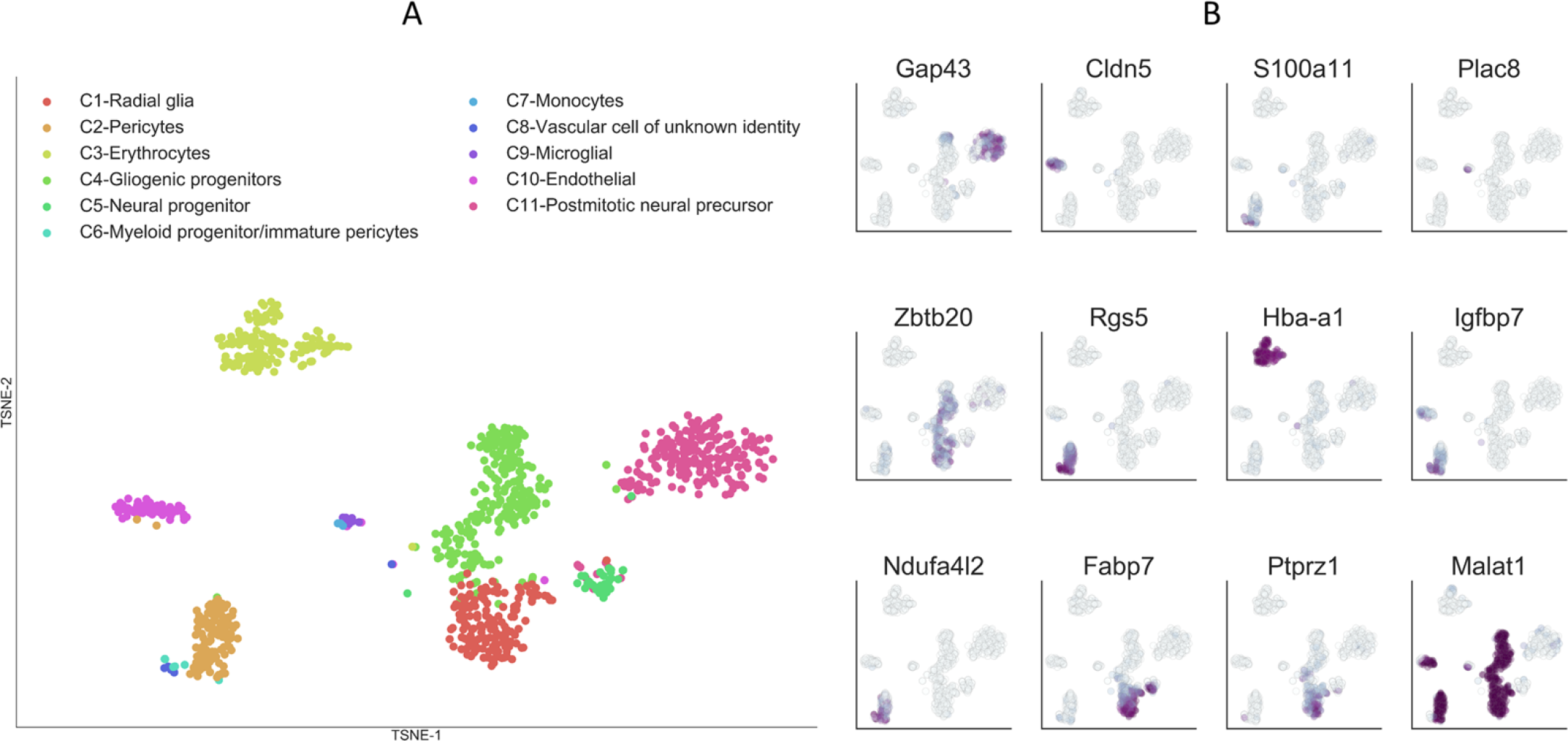
Single-cell clusters of mouse embryonic hypothalamus identified by PanoView. (A) Visualization of identified cell types from embryonic hypothalamus using TSNE. (B) Relative expression of 12 selected marker genes in embryonic hypothalamus.

### Summary

In this study, we have described the development and performance of PanoView to identify cell subpopulations in single-cell gene expression datasets. Without any tuning of the parameters, PanoView produced reasonable clustering results in 1,200 simulated data and 11 published scRNA-seq datasets. Furthermore, without any adjustment, PanoView was able to identify rare cell types in both simulated data and scRNA-seq datasets. The robust performance of PanoView may be the result of both searching cell clusters one by one in the evolving PCA space and improved density-based clustering. Note that it is possible that other clustering methods may show improved performance once we fine-tune their parameters with the input from experienced experts. We believe that PanoView can offer reliable performance with moderate computational cost and can be applied to diverse types of scRNA-seq dataset. The clustering of single cells will automatically identify cell specificity. After the identification of cell types, we are also able to determine the marker genes that show specific expression in each cell type (e.g. Fig. 6B). We believe that the cell atlas and the corresponding marker genes will be a valuable resource to study various biological processes.

## Materials and methods

### PanoView algorithm

The key of PanoView is to iteratively search clusters in different sets of variable genes. Our algorithm first performs PCA reduction based on a set of variable genes (defined below). By choosing the first three principal components which explain the largest variance across all cells, PanoView then applies a novel density-based clustering approach, ordering local maximum by convex hull (OLMC), to cluster cells into multiple groups. These groups are evaluated by their variances and the Gini index in the current gene space. PanoView then identifies the best “mature” cluster that is the one with the lowest variance, and the rest of the cells will be put into the next iteration. A new set of variable genes is determined with the remaining cells and the same procedure (PCA reduction and OLMC) is repeated. The iteration of PanoView is terminated when no more cluster can be produced, or Gini index reaches a threshold. Next, PanoView produces a hierarchal dendrogram for all generated clusters and merges similar clusters based on the cluster-to-cluster distance.

A pseudo-code is provided as the following to detail as to how PanoView works:

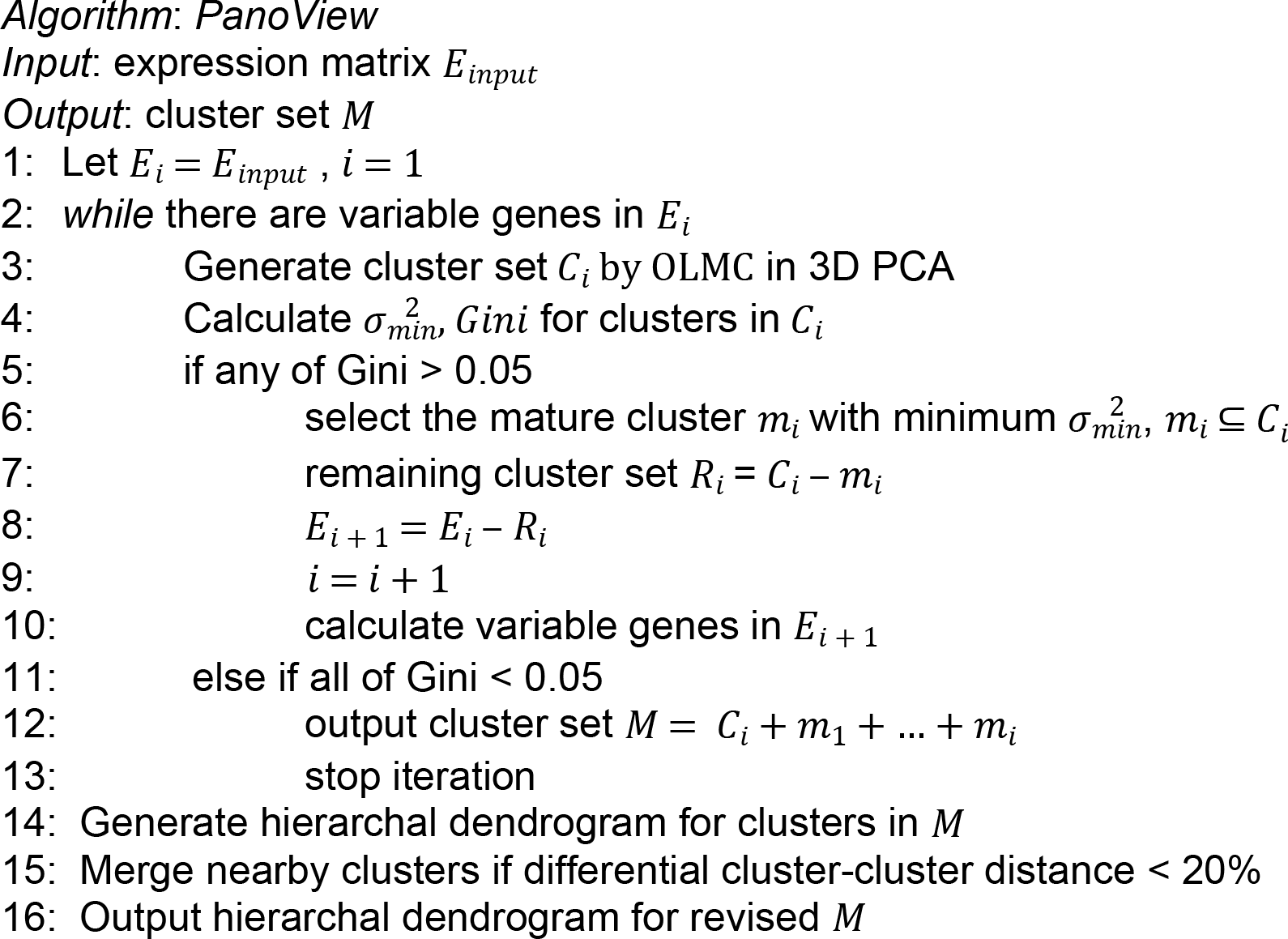

### Variable genes for PCA

We adopt the procedure described in Macosko *et al* to find variable genes(3). First, all genes are grouped into 20 bins based on their average expression levels. Second, the ratio of variance and mean for genes in each bin is calculated. Third, z-normalization is performed using the ratio of variance and mean in each bin and using the z-score as a threshold to obtain a set of variable genes. The default value of z-score is 1.5 (Zscore=1.5). We also exclude the lower expressed genes whose average expression is less than 0.5 (Genelow=0.5). This selection of variable genes is carried out during each iteration of PanoView.

### Ordering Local Maximum by Convex Hull (OLMC)

For clustering single cells, we developed ordering local maximum by convex hull (OLMC), a density-based clustering, to identify local maximums in three-dimensional gene space. First, we compute the pairwise Euclidean distance of cells. The distances were grouped into *B*_*c*_ bins (default value = 20) with equally distance interval. The *R*_*c*_ is the bin interval of the histogram that represents the calculated distribution based on the input dataset. Second, we applied the k-nearest neighbors algorithm implemented in Scikit(29) to compute the number of neighbors within distance *R*_*c*_ for each cell. The cells are then ordered based on the number of neighbors, with each cell annotated as *P*_*i*_, where *i* is the ranking index from 1 to the total number of cells. *P*_1_ represents the global maximum in the space. Third, the cells are equally grouped into *B*_*g*_ bins based on the distance to *P*_1_. The cells in the first bin are considered as the first group *G*_1_, and a convex hull *H*_1_ that compose of a set of vertices is constructed. Third, we search for the next local maximum density. Assuming *P*_*m*_ is the first one from the remaining ranked cells, we first define 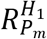 as the distance to the nearest vertices of *H*_1_ and 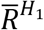 is the average of pairwise distance for the vertices of convex hull *H*_1_. If 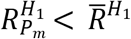, *P*_*m*_ will be added into the group *G*_1_, and corresponding convex hull *H*_1_ is updated (i.e. expanding), suggesting *P_*m*_* is not a local maximum. If 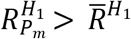, the new local maximum *P*_*m*_ is located and the corresponding convex hull *H*_2_ is constructed based on the distance to *P*_*m*_. The searching for the next local maximum would be ended if the number of remaining cells is not sufficient to construct a convex hull. Once every local maximum density is located, assign every cell to the nearest local maximum densities. To sum up, the key of OLMC algorithm is to first find where the global maximum density is and use convex hull to locate the next local maximum.

To illustrate OLMC, a toy model consisting of 500 random points is provided (Fig 1B-D). In Fig 1B, each number represents the number of neighbors within *R*_*c*_ = 0.5. The histograms in 1C represent the distance to local maximums and are built by *B*_*g*_ = 10. The number of 27 in Fig 1B is where the highest density is. The first convex hull (the cyan in Fig 1D) is constructed by the points within the first bar (Fig 1C) of the distance histogram. After removing the points in the cyan convex hull, the next point with the highest density is where number of 23 is, and the second convex hull is constructed by the points in the first bar (in green) of the second histogram that is calculated by distance distribution to the point of 23, a local maximum density. Followed by the same procedure, the next local maximum (point of 22 in yellow) is located and the third convex hull is built. In the end, OLMC identifies the locations of three local maximums, and assign rest of the points to the nearest local maximums.

In PanoView, the goal is to find as many clusters as possible during the iterations. Therefore, we adopted a heuristic approach to optimize the bin size *B*_*g*_ that controls the histogram of distance to local maximums for constructing convex hulls. We generated a simulated data of 500 2D points to illustrate the optimization (S3 Fig). By incrementally increase the bin size by 5, OLMC would reach a saturated state that no more local maximums can be located. We carry out the optimization until the saturated state or the bin size of 100 (Maxbb = 20)

Due to the computational efficiency, this optimization is only activated when the number of cells during iterations is smaller than CellNumber=1000. Otherwise, the default *B*_*g*_=20.

### Cluster evaluation in PanoView

One crucial step in PanoView is to evaluate the clusters produced by OLMC for locating the “mature” cluster during each iteration. The idea is to use Gini index to evaluate the inequality of clusters. PanoView first calculates the pairwise correlation distance *X*_*i,j*_ for every cell within each cluster using 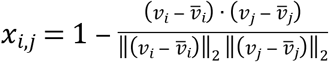
where 𝑣_*i*_, 𝑣_*j*_ are *n*-dimensional vectors and 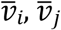 are the means of the elements of vector 𝑣_*i*_, 𝑣_*j*_, respectively (30). The algorithm then calculates the variance *σ*^2^ of distances *x*_*i,j*_ for each cluster and ranked the clusters in the descending order.

PanoView then calculates the Gini index *G*_*i*_ (*i*=2, to *n*), for the top *i* clusters. Here n is the total number of clusters in this iteration. The Gini index(31) was defined as

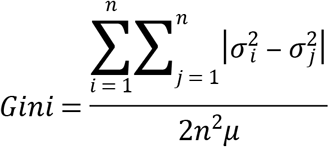

where 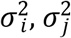 are the variances in a population of variances, *n* is the number of variances, and *μ* is the mean of a population of variances.

If there is a Gini smaller than the threshold of 0.05, PanoView will keep the cluster with the minimum variance (i.e. the “mature” cluster) and put the rest of cells into the next iteration.

### Generation of simulated datasets

We used Scikit’s sample generator (29) with default parameters except the number of clusters and standard deviation within each cluster. These datasets served as the ground truths to evaluate the ability to identify cell subpopulations for chosen computational methods. Each simulated dataset consists of 500 cells and 20,000 genes, with expression values in the range of 0 to 10,000. The cells are equally divided into numbers of clusters based on randomly generated *n* centers (3 ≤ *n* ≤ 22). In each cluster, cells are dispersed around the center of the cluster with a given standard deviation (*SD* = 0.5,1,2). For each *n*, we generated 20 random configurations (i.e. datasets). In total, we generated 1,200 different random datasets.

For evaluating the ability to identify rare cell-types, we followed the same procedure to generate simulated datasets. The number of clusters ranged from 3 to 15, and the standard derivation of each cluster was 1. In each dataset, we randomly picked one cluster and removed 90% of the cells from that cluster. This cluster was defined as the rare cell subpopulation. In other words, the size of the rare cluster is about 0.6% to 3% of the total population. We also varied the random state of the generator by 20 random numbers to have a total of 260 random datasets. The command line to generate the simulated datasets in python’s Scikit is “make_blobs(*n_samples*=*500*, *n_features*=*20000*, *centers*=*None*, *cluster_std*=*1.0*, *center_box*=*(−10.0*, *10.0)*, *shuffle*=*True*, *random_state*=*None*)”. The python code for generating simulated datasets are available at PanoView’s Github repository.

### Real single-cell RNA-seq datasets

We used the following 11 scRNA-seq datasets in our study. Yan et al profiled transcriptomes of human preimplantation embryos and human embryonic at different passages (32) (GSE36552). Goolam et al profiled transcriptomes of mouse preimplantation development from zygote to late blastocyst (33) (E-MTAB-3321). Deng et al used scRNA-seq to study the allelic expression of mouse preimplantation embryos of mixed background (CAST/EiJ × C57BL/6J) from zygote to late blastocyst (34) (GSE45719). Pollen et al used low-coverage scRNA-seq to study the development of the cerebral cortex in hiPSCs (35) (SRP041736). Patel et al reported expression profiles of single glioblastoma cells from 5 individual tumors (6) (GSE57872). Usoskin et al used single-cell transcriptome analysis to study cell types of mouse neurons (36) (GSE59739). Villani used scRNA-seq to classify dendritic and monocyte populations from human blood (37) (GSE94820). Zeisel used scRNA-seq to study the transcriptome of mouse somatosensory cortex S1 and hippocampus CA1(2) (GSE60361). Tirosh et al used scRNA-seq to study genotypic and phenotypic states of melanoma tumors from 19 patients (38) (GSE72056, GSE77940). Baron et al used inDrop technique to profile the transcriptomes of human and mouse pancreatic cells (5) (GSE84133). Campbell et al used Drop-seq to study transcriptomes of mouse arcuate nucleus and median eminence (39)(GSE93374).

### Benchmark with other clustering methods

For parameters in pcaReduce, we used the default setup (*nbt* = 1*, q* = 30*, method* = “s”). For key parameters in Seurat, we used *model.use* = “negbinom”, *pcs.compute* = 30, *weight.by.var* = FALSE, *dims.use* = 1:10, *do.fast* = T, *reduction.type* = “pca”, *dims.use* = 1:10. For key parameters in SCANPY, we used *counts_per_cell_after*=1e4, *min_mean*=0.0125, *max_mean*=3, *min_disp*=0.5, *max_value*=10, *n_neighbors*=10, *n_pcs*=40. For RCA, we used the default setup. For SC3, we used the default setup and *sc3_estimate_k* as the final clustering output. In the Baron dataset, SC3 only reported the clustering result for 5,000 random cells due to the activation of SVM. We had to use these reported 5,000 cells to calculate the ARI value. For DBSCAN, we first did PCA reduction with Scikit’s default setup and adjusted *epsilon* and *minPts* based on the visualization of PCA space. We also used Scikit’s default setup for executing Kmeans (*n_clusters* = k, *init* = ‘random’) and TSNE (*n_components* = 2, *random_state* = 1, init = ‘random’, *n_iter* = 1000).

For benchmarking RaceID2 in our simulated datasets, we used the default setup from the manual and did not pass the step of *findoutliers*. Therefore, we used @*cluster$kpart* as the final clustering result. For benchmarking GiniClust in our simulated datasets, we used the default parameters from the manual except for *Gini.pvalue_cutoff*. We adjusted it from 0.0001 to 0.005 because the default value of 0.0001 did not produce useable clustering results.

### Evaluation of performance in detecting rare cell types

We used recovery rate and false positive rate to evaluate the performance of clustering methods on detecting rare cell types. In each simulated dataset, we always have one rare cell cluster and *n* (*n*=2 to 14) major cell clusters. If the rare cell cluster was perfectly detected with the correct number of cells within the cluster, we considered that the algorithms recovered the rare cell type. On the other hand, if cells from a major cluster were grouped into multiple clusters and at least one of the sub-cluster had the size less than 10% of the major cluster, we considered that the algorithm generated a false positive rare cell type.

### Animals

Timed pregnant mice (Charles River Laboratories, MA, USA) were housed in a climate-controlled pathogen-free facility, on a 14 hour-10 hour light/dark cycle (07:00 lights on-19:00 lights off). All experimental procedures were pre-approved by the Institutional Animal Care and Use Committee of the Johns Hopkins University School of Medicine.

### Single-cell RNA-Seq library generation and analysis

Hypothalamic tissue dissected from embryonic day (E) 16.5 C57BL/6 mouse embryos under a dissecting microscope in cold 1x HBSS (Thermo Fisher Scientific, MA, USA). A total of 6 embryos were dissected. Dissected tissues were incubated in papain solution (Worthington, NJ, USA) at 37’C for 15 minutes. Papain activity was stopped as following manufacturer’s protocol, and fire-polished Pasteur pipette was used to gently pipette tissues up and down to dissociate tissues into single cells. Dissociated cells were filtered through 40 uM strainer and washed once in Neurobasal media (Thermo Fisher Scientific), and cells were resuspended in Neurobasal media with 1% bovine serum albumin. Approximately 17,000 live cells were loaded per sample in order to capture transcripts from roughly 10,000 cells. Estimations of cellular concentration and live cells in suspension was made through Trypan Blue staining and use of the Countess II cell counter (ThermoFisher). Single cell RNA capture and library preparations were performed according to manufacturer’s instructions. using 10x Genomics Chromium Single Cell system (10x Genomics, CA, USA) using the v1 chemistry, following manufacturer’s instructions and sequenced on Illumina MiSeq system (Illumina, CA, USA). Sequencing results were processed through the Cell Ranger pipeline (10x Genomics) with default parameters to generate count matrices for subsequent analysis. The total number of single cells is 959, and the total number of reads is 15,365,879. The mean reads per cell is 16,022, and total genes detected is 15,223. The median number of genes per cell is 617.

### Software availability

PanoView is available as a Python module at https://github.com/mhu10/scPanoView. To run a clustering analysis with default parameters in PanoView, we run two command lines, RunSearching(GeneLow = ‘default’, Zscore = ‘default’) and OutputResult(fclust_height = ‘default’). The complete user manual is provided at Github repository.

## Supporting information captions

S1 Fig: **The clustering results of different parameters in DBSCAN**. Clustering results of DBSCAN with different sets of parameters (*epsilon* and *minPts*). ARI value represents the similarity between the DBSCAN result and the ground truth. The value of 1 would indicate the clustering membership is the same as the ground truth.

S2 Fig: **The similarity between the results of DBSCAN with different parameters**. The pairwise comparison of clustering results from Figure S1. Each value represents the ARI of the results from two different parameter sets.

S3 Fig: **The heuristic approach for estimating bin size in OLMC**. (A) The result of OLMC on 500 random 2D points analyzed using different bin sizes. (B): Optimal bin size is between 20 to 45 for this simulated data.

S4 Fig. **The results of different clustering methods in Patel dataset**. Comparison of different clustering methods using the Patel dataset. Visualization of clusters was generated by TSNE. Panel A shows the original clustering results from the Patel et al publication. Panels B-F show the clustering results with different methods.

S1 Table: **Key parameters in some computational methods for scRNA-seq**. Key parameters for some computational methods used in scRNA-seq

S2 Table: **scRNA-seq datasets used in this study. Published scRNA-seq datasets used in this study**. N is the total number of cells. K is the reported number of clusters in the original published studies

S3 Table: **The clustering result of different computational methods in published scRNA-seq datasets**. The result of ARI calculation in scRNA-seq datasets

S4 Table: **The clustering result of PanoView with 500 random parameter sets in published scRNA-seq datasets**. 500 clustering results of PanoView that includes ARI, values of parameters, and the computational cost.

